# Characterization of the Peer Review Network at the Center for Scientific Review, National Institutes of Health

**DOI:** 10.1101/007377

**Authors:** Kevin W. Boyack, Mei-Ching Chen, George Chacko

## Abstract

The National Institutes of Health (NIH) is the largest source of funding for biomedical research in the world. This funding is largely effected through a competitive grants process. Each year the Center for Scientific Review (CSR) at NIH manages the evaluation, by peer review, of more than 55,000 grant applications. A relevant management question is how this scientific evaluation system, supported by finite resources, could be continuously evaluated and improved for maximal benefit to the scientific community and the taxpaying public. Towards this purpose, we have created the first system-level description of peer review at CSR by applying text analysis, bibliometric, and graph visualization techniques to administrative records. We identify otherwise latent relationships across scientific clusters, which in turn suggest opportunities for structural reorganization of the system based on expert evaluation. Such studies support the creation of monitoring tools and provide transparency and knowledge to stakeholders.

## Introduction

The National Institutes of Health (NIH) is the premier biomedical research agency in the United States. NIH supports both basic and applied biomedical research largely through awards of grants to extramural applicants. The principal basis for award is scientific merit as determined by peer review.

In 1946, the first study section was assembled at NIH to conduct peer review of applications for support of research on syphilis [1]. Also known as a Scientific Review Group (SRG), a study section is a panel of expert scientists assembled to evaluate a set of grant applications. SRGs are of two principal types: chartered SRGs and Special Emphasis Panels (SEPs). Chartered SRGs have defined scientific interests, meet three times a year, and have relatively stable membership while SEPs are typically assembled for a single meeting.

Peer review at NIH has evolved significantly since the first study section meeting. Within NIH, the Center for Scientific Review (CSR) manages the peer review process for the large majority of grant applications received. In 2014, sixty-eight years after the first study section meeting, more than 170 chartered SRGs exist at CSR, each centered on a scientific theme, e.g., the Nuclear and Cytoplasmic Structure/Function and Dynamics (NCSD) study section. SRGs at CSR are clustered into 25 Integrated Review Groups (IRGs), each again centered on a scientific theme of broader scope than an SRG (see Supplementary Information). The scientific interests of the IRGs at CSR are organized, somewhat heterogeneously, along sub-disciplines, organ systems, basic versus applied research, and disease-specific interests, e.g., Cell Biology (CB), Oncology 1-Basic and Translational (OBT), and Digestive, Kidney, and Urological Systems (DKUS). The IRGs are further aggregated into five Review Divisions, which vary in focus from basic to applied research.

Implicit in the design of this organizational structure are organizational objectives: that the system, as a whole, should stay abreast of scientific trends and provide adequate coverage, enable fair competition, accommodate workloads and their temporal fluctuations, exhibit transparency, and enjoy public confidence while being austere in consuming resources. The scale and scientific impact of these operations is very large; in fiscal year 2012, reviews for 56,000 grant applications were managed by CSR, resulting in new grant awards of roughly $3 billion (rounded to the nearest 1000 applications and $100 million). Thus, it is critically important that the system of peer review performs competently.

A major goal of this study was to provide a comprehensive characterization of the current structure of CSR?s peer review system from a scientific perspective as a first step towards identifying avenues for improvement. Technological advances have enabled us to apply textual analysis and bibliometric techniques to administrative records, text from grant applications, and large-scale bibliographic data to generate analytic visualizations that provide a novel system-level description of the scientific structure of CSR?s study section network that can be used as a model in evaluation studies.

A variety of techniques have been used over the years to identify and visualize the implicit structure of various domains from information associated with those domains. Common to all of these techniques is a generic process flow [2] in which units of analysis are chosen, similarity between those units is calculated, and the resulting similarity matrix is used to generate a view of the domain, often by visualizing it in the form of a network.

A majority of the work done in this area has been to visualize the structure of science or of particular scientific domains by applying the above process to bibliographic data from the Web of Science, Scopus, PubMed, or other literature databases. Typical units of analysis include articles [3], authors [4], keywords, journals [5,6], and subject categories [7]. Grants [8], patents, and patent categories [9,10] have also been mapped. Similarity between objects (or units) is typically calculated using co-occurrence in the feature space associated with the objects; co-citation [11] and co-word [12] are prominent examples of techniques based on co-occurrence. More complex ways of calculating similarity, such as the well-known vector space model (e.g., Salton’s cosine) [13], are also often used to calculate pairwise similarities from a matrix of co-occurrence values.

Once similarity values between objects are calculated, a variety of methods can be used to generate a layout (or visual map) of those objects. One class of algorithms, known as graph layout algorithms [14,15], considers each similarity to be a weighted edge between objects, and uses these edges to create a graph-based visualization of the relationships between the objects. Other techniques, such as multidimensional scaling, attempt to create an optimum layout of objects using distances (or dissimilarities) between them rather than operating on the network of similarity-weighted edges.

To achieve the goal of visualizing and characterizing the peer review network, chartered SRGs are designated as our unit of analysis in this study. We restricted our analysis to SRGs from 24 of 25 IRGs (Fig. 1), excluding one IRG that was not considered representative on account of containing only one SRG that was atypical. Similarity values between SRGs were calculated using a number of different data types and features associated with the study sections. Finally, the relationships between the SRGs were visualized as networks using calculated similarity values.

**Figure 1.**
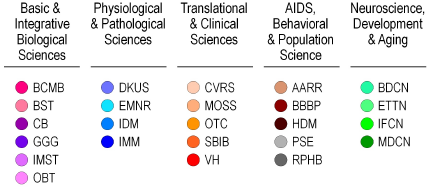
Listing of IRGs at CSR by Review Division. Each IRG has been assigned a color to be used in subsequent figures. Divisions are grouped by color. SRG to IRG assignments, acronyms and full names are provided in Supplementary Information.

## Results and Discussion

The organizational structure of CSR lends itself to representation as a graph network where each SRG is a node and edges can be drawn to other nodes on the basis of scientific relatedness. We analyzed administrative records corresponding to 72,526 applications, 42,564 applicants, and 11,896 unique reviewers from the fiscal years of 2011 and 2012 (see Materials and Methods). We calculated and visualized several different versions of the SRG network as a part of this study. Six different representations of the full SRG network were created using different measures of relatedness between study sections. A consensus network that takes all of the different networks into account was also created.

Figure 1 shows the high level administrative structure of the CSR peer review system. As a first step towards evaluating whether this system satisfies the organizational objectives described in the Introduction, we need to understand how the scientific structure of the system, as defined by similarities between the topics embodied by each IRG and SRG, differs from the management perspective. Large differences might warrant an evaluation by expert scientists to consider changes to the organizational structure.

The process for depicting the structure of each SRG network consisted of two major steps. First, the relatedness of pairs of study sections was determined. Relatedness was defined in six different ways within three main groups of feature characteristics, as follows:

- NIH classification
  - Research, Condition, and Disease Categorization-RCDC (http://report.nih.gov/rcdc/) profiles
- Data associated with reviewers (publication years 1996-2011)
  - Publications authored by reviewers (REV-P)
  - Titles and abstracts of publications authored by reviewers (REV-T)
  - Cross-citation patterns between reviewers (REV-C)
- Data associated with applications and applicants
  - Text (title, abstract, specific aims) of grant applications (APP-T)
  - Applicant publications (APP-P)

Details on the calculation of each of these six types of relatedness are given below in the Methods and Materials: SRG Relatedness section. Each relatedness matrix can be represented as a directed graph where the study sections are vertices (or nodes) and the relatedness values are weighted edges that connect these vertices. Once the similarity matrices are calculated, a graph layout algorithm is used to create a visual picture of each graph.

Using the feature data mentioned above, we found that there were very few pairs of study sections with a zero-valued similarity; a weighted edge exists between nearly all pairs of study sections. Sparse networks are far easier to interpret than non-sparse networks. Accordingly, we desired to create diagrams based on the dominant network rather than on the full network. Initial studies showed that restricting edges to the top-3 similarity values per study section sufficed to generate a connected graph that exhibited the dominant characteristics of the network. In addition, previous work has shown that layouts based on only the strongest few edges per object form more accurate clusters of objects than layouts based on large numbers of edges [16]. Thus, the layout was based on the top-3 similarity values per study section rather than on the full set of similarities. Each layout was created with Pajek [17], using the Kamada-Kawai layout algorithm [15] since it produces very readable graphs for networks of modest size (200 nodes). When creating the visualizations, each IRG was designated using a different color (defined in Figure 1), and links between study sections within the same IRG were shown with that color. Links between SRGs that are not in the same IRG are in black. In addition, the directions of the edges are shown using arrows. Arrows point from the choosing study section to the chosen study section. Links with arrows at both ends denote that each SRG in the pair was in the top 3 list of its corresponding SRG.

Figure 2 shows the six visualized networks along with the IRG color scheme. To facilitate comparison, all six networks have been rotated to have a similar orientation (BCMB [red] at the upper left and MDCN [dark green] at the upper right).

**Figure 2.**
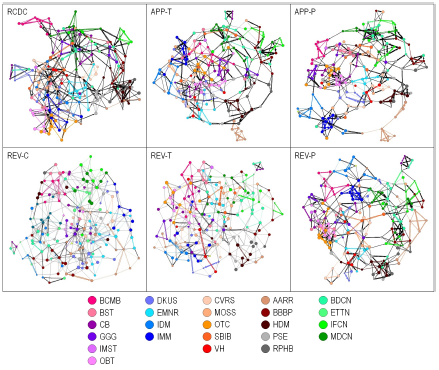
Study section networks generated using six different features associated with SRGs. Edges between SRGs within the same IRG are colored with the IRG color. Edges between study sections in different IRGs are colored black. Arrows point from the choosing study section to the chosen study section. Links with arrows on both ends denote that each study section is in the top 3 list of the paired study section.

These six different networks reveal several features. First, each network comprises a single component, i.e., no node is disconnected. Second, there are many areas of color concentration in all six networks, which suggests that all six measures, while they are based on different features, reconstruct the existing IRG structure to a relatively high degree. This point was further investigated by calculating the percentage of edges in each network that are within an IRG. Nearly half of all edges were within IRGs, as shown in Table 1.

In contrast, the fraction of within-IRG edges in a random network was only 4%, showing that all six networks were far from random. Table 1 also shows the number of bidirectional edges, which is greater than half in most cases, and very small in the random case. Bidirectional edges indicate that both members of a pair of SRGs consider the other to be within its top-3 and large numbers of bidirectional edges are consistent with the concept of a robust network. In addition, color groups (e.g., blues, greens, oranges) denoting IRGs that are a part of the same Review Division are largely in local subsections of the maps. For example, the browns and grays (AIDS, Behavioral & Population Science Division) are found together and at the right side of five of the six maps in Figure 2. This suggests that the grouping of IRGs into Review Divisions, although organized under a different and more coarsely granular principle than the networks shown here, is roughly consistent with the science-based groupings of Figure 2. The six different networks also show substantial connections between IRGs. This is not surprising given that specialties in the life sciences are known to be highly interlinked [18], but this observation also identifies areas that may merit study towards the organizational interests described in the Introduction.

**Table 1.** Comparison of within-IRG links for each study section network.

| Method | #SRG | #Links | %Links<br>within IRG | # Links<br>Bidirectional |
| --- | --- | --- | --- | --- |
| Random* | 174 | 522 | 4.1%(0.6%) | 8.6 (3.2) |
| RCDC | 174 | 522 | 50.8% | 244 |
| REV-P | 180 | 540 | 46.7% | 304 |
| REV-T | 180 | 540 | 47.6% | 318 |
| REV-C | 180 | 540 | 41.9% | 238 |
| APP-T | 180 | 540 | 45.4% | 314 |
| APP-P | 180 | 540 | 47.0% | 314 |
| Consensus | 174 | 290 | 57.9% |  |

\*Averages (standard deviations) from 20 trials.

The six networks of Figure 2 are each based on a different set of criteria in determining relatedness between SRGs. There is no *a priori* way to say which of them is the most valid-they are simply different representations. Rather than choosing one of these criteria to use as the basis for representing the SRG network, we propose that a consensus map of the study section network would likely be a more accurate representation of the actual network.

A consensus map is not an ‘average map’, but rather one in which the links between nodes are highly rated using a set of rules that consider all of the inputs [19]. The six different measurements detailed above were created using three different sets of criteria: one was created using the RCDC fingerprints for each study section, three were created using various representations of reviewer expertise, and two were created using different representations of application content. All of these measurements were considered as input to the consensus map.

To generate a consensus map, we first had to identify those links between SRGs that were highly ranked by multiple criteria (consensus links). The following protocol was used to identify consensus links (Table 2), and then to generate a consensus study section network. It is difficult to combine relatedness measures based on different approaches that have different range values. Thus, this calculation uses rank orders rather than relatedness values. The six input metrics mentioned above represent three sets of criteria. We decided that each of these three criteria (i.e., RCDC, reviewers, applications/applicants) should contribute equally to the consensus ranking for each study section pair, i.e., SRG1 and SRG2 and used the following protocol.

1. A single rank value was calculated for each SRG pair based on reviewer data as the highest rank value (best = 1) for that pair among the three different reviewer-based relatedness metrics (REV-P, REV-T, REV-C), i.e., from left to right in the Reviewer column of Table 2. The highest rank value among the three is denoted in the parentheses.
2. A single rank value was calculated for each SRG pair based on application/applicant data as the highest rank value (best = 1) for that pair between the two relatedness metrics (APP-T, APP-P), i.e., from left to right in the Application/Applicant column. The highest rank value among the two is denoted in the parentheses.
3. The rank value for each SRG pair from the RCDC similarity metrics was used without comparing it to any other metric, i.e., RCDC column.
4. Using these three rank values, the worst rank value for each SRG pair was selected as the consensus rank value for that pair, resulting in the Consensus column. A set of example rankings and the resulting consensus rank are shown in Table 2. Note that the initial steps used the highest rank for each SRG pair for each of the three major criteria, while this consensus-finding step used the lowest ranking among the three sets of criteria. The use of the highest ranking within a criteria set is based on the logic that each set should give an SRG pair its best chance of being included in the final consensus set. This is balanced by the use of the lower ranking in the consensus-finding step.
5. Given that we now had the full list of study section pairs with their consensus rank values, we needed to decide which of those pairs (edges) to use to create the consensus network. A consensus network should include all SRGs while using a minimum number of edges. In addition, only the best edges (those with the highest consensus rank values) should be used. Table 3 shows the numbers of edges and SRG coverage as a function of consensus rank values. If all edges with rank ≤ 3 are kept, most of the study sections (166/174) are included. At rank ≤ 4, only three more study sections are included, while 88 edges are added, increasing the complexity of the network.
6. Since simple networks are far easier to understand than more complex networks, we decided to use the network based on rank ≤ 3, and augment it with the highest ranked link for the eight SRGs that were missing. This has the effect of adding these SRGs into the network while only adding 16 links.
7. We then visualized the resulting network. To create a layout, rank values were converted to similarities as Sim = 5 - Rank, with a minimum Sim value of 1.0. The resulting similarity file was used as input to the Kamada-Kawai layout algorithm in Pajek. This layout was further modified by hand to reduce edge crossings and to create additional space between study sections for greater legibility.

**Table 2.** Examples of consensus rank calculations.

| SRG1 | SRG2 | RCDC | Reviewer | Application | Consensus |
| --- | --- | --- | --- | --- | --- |
| ACE | BSCH | (2) | 3 4 3 (3) | 1 1 (1) | 3 |
| ACE | CRFS | (4) | 10 2 4 (2) | 7 8 (7) | 7 |
| ADDT | AMCB | (2) | 4 6 3 (3) | 1 1 (1) | 3 |
| ADDT | AIP | (4) | 2 1 1 (1) | 2 2 (2) | 4 |
| AED | DPVS | (1) | 1 2 1 (1) | 1 2 (1) | 1 |
| AED | BDPE | (5) | 3 3 3 (3) | 3 4 (3) | 5 |
| AICS | VCMB | (2) | 1 1 1 (1) | 3 3 (3) | 3 |
| AICS | CICS | (4) | 7 5 5 (5) | 7 4 (4) | 5 |

Separate ranking values for each approach are shown in the appropriate columns. Reviewer (REV-P, REV-T, REV-C ), Application (APP-T, APP-P ). Single rank values for each set of criteria are shown in parentheses.

**Table 3.** Numbers of study section pairs by consensus rank

| Rank | #Links | #SRGs | %Links/SRG |
| --- | --- | --- | --- |
| 1 | 79 | 104 | 0.76 |
| < 2 | 169 | 149 | 1.13 |
| < 3 | 274 | 166 | 1.65 |
| < 4 | 362 | 169 | 2.14 |
| < 5 | 466 | 172 | 2.71 |

We also measured the contributions of the six networks (Fig. 2) to the consensus network. Of 209 consensus edges, 202 (97%) were found in the RCDC network, 184 (88%) in APP-T, 177 (85%) in APP-P, 171 (82%) in REV-P, and 146 (70%) in both the REV-T and REV-C networks.

A visual map of the consensus SRG network is shown in Figure 3. (This consensus network is also provided with higher-order labeling as Figure 4.) The network consists of one large component with 162 SRGs, and three small components containing 12 SRGs. The three small components are at the top, upper right, and lower right of the map, and have been linked to the large component using dashed lines based on their strongest links to SRGs in the large component. As with the input maps shown in Figure 2 (note that the maps of Figure 2 are oriented to roughly match that of the map in Figure 3), study sections in the same IRG are often connected and proximal to each other in the map. For example, most of the SRGs in the BCMB IRG are adjacent to one another (red, upper left).

**Figure 3.**
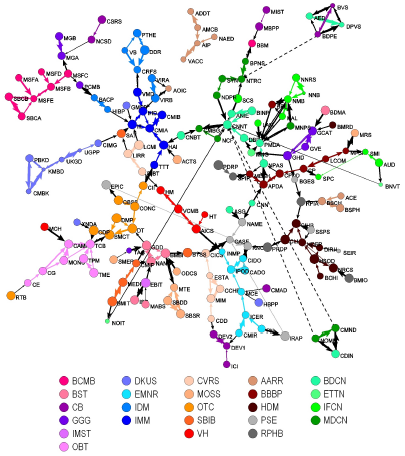
Consensus study section network. Study section names for each acronym are available in Appendix Table 2. Edge width corresponds to rank the thickest edges represent *rank* = 1, etc. Edges between study sections in the same IRG are colored with the IRG color. Edges between study sections in different IRGs are colored black. Dashed lines indicate where the smaller components connect to the main component, but with lower ranked links.

**Figure 4.**
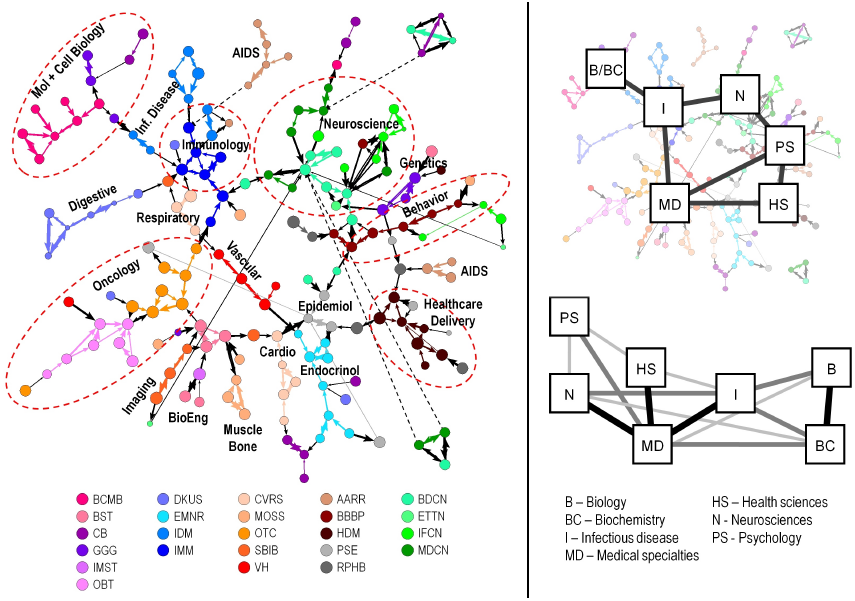
Left. Consensus study section network with higher order labels, which were created manually based on names of the SRGs and IRGs in different regions. This network is identical to Figure 3. Dashed ovals do not denote fixed scientific boundaries, but are intended to approximate them. Right: Comparison of the high-level structure inferred by the consensus study section network with the biomedical portion of the K/B consensus map of science. Discipline names are noted.

The map also provides a context in which to consider different ways to group study sections into IRGs, and to group IRGs into Review Divisions. For example, the AARR IRG is split, with five of its SRGs (tan) at the upper middle of the map, and three study sections at the middle right of the map, and one connected with another component. While our analysis suggests an opportunity for rearrangement, the existing clusters can be explained by an administrative requirement for clustering and expedited review of AIDS applications [20]. This example illustrates the interplay between administrative and scientific objectives that contribute to the structure of the peer review network at CSR. Another case is the small component at the upper right, which consists of two SRGs from BDCN (light green) and two study sections from CB (purple). Not only are these SRGs from different IRGs, but they are also from IRGs that are in different Review Divisions. All four SRGs are organized along different themes but evaluate applications concerned with biology of the visual system.

Most of the SRGs from three IRGs in the Division of Physiological and Pathological Sciences (DKUS, IDM, IMM) are connected to each other at the upper left of the map, while the fourth IRG (EMNR) is at the bottom of the map is widely separated from the other IRGs. This IRG, EMNR, seems to be more closely situated with IRGs that are colored orange, most of which are at the bottom and bottom left of the map. Greens are typically found in the upper right portion of the map, while the brown/black hues are at the far right. IRGs with purple hues are perhaps the least coherent set on the map; these can be found at the right (GGG), top (CB), upper left (BCMB) and lower left (OBT, BST). All five are concerned with basic research and technology development. We speculate that the distributed nature of the nodes from the basic science areas may reflect diffusion of knowledge and technology transfer into more applied fields as well as inspiration in the reverse direction. Elucidating these possibilities are topics for future study. In general, however, the groupings of IRGs on the consensus map (Figure 3) reflect the groupings of IRGs in the management structure (Figure 1) to a remarkable degree, particularly in light of the fact that the current structure has evolved in response to a combination of scientific, administrative, and societal inputs.

It is also instructive to compare the consensus SRG structure with existing maps of science. Figure 4 (right side) compares the consensus SRG map coded using the high-level disciplinary structure used in the Klavans/Boyack (K/B) consensus map of science [19] with the biomedical portions of that map. The majority of the strong links in the K/B consensus map are preserved in the consensus SRG map, suggesting that the consensus SRG map based on few links is consistent with an accepted high-level structure of science, while recognizing that resolution is lost with higher-level descriptions.

In evaluating the contributions of the six individual networks (Fig. 2) we note that the RCDC network alone contains information to reproduce 97% of the edges in the consensus network. Thus, the RCDC network alone may be adequate to construct the network but with some loss of resolution due to the inclusion of a similar number of non-consensus edges. In our study, we observe that the networks based on features of applications and applicants are more closely related to the consensus than are the networks based on features of the reviewers.

Overall, this study identifies a network of SRGs related by scientific interests that are partially coincident with the current administrative structure, an evolutionary product of a peer review system subject to both scientific and management constraints. Our representation of the network is a model that supports evaluation of the existing system enabling the design of improvements against defined optimality criteria based on organizational objectives (described in the Introduction). Given the complexity and nature of constraints on the system, refinement of the network for greater utility to its stakeholders must simultaneously consider clustering (grouping of SRGs into IRGs), classification (assignment of applications and reviewers to SRGs), and balance (workloads) while being considerate of the preferences of applicants. In addition, administrative constraints such as the number of SRGs that could be supported given available resources should be applied to synergize with scientific objectives. Although mathematical techniques can be brought to bear on these questions, answers ultimately require human intervention, particularly given that several optimal solutions could be identified. Requiring that any SRG should have application numbers within a specified range would satisfy the organizational interest in fair competition. Designing each SRG for optimal breadth and depth of science to enable such competition, and designing the system to provide coverage for the spectrum of scientific disciplines represented by the input set of applications each year are questions that will be the subject of further studies.

## Materials and Methods

### Data

Datasets of grant applications, applicants, and reviewer information for SRGs organized by CSR in the fiscal years of 2011 and 2012 were assembled by querying IMPAC II, the NIH database of grants information. The application dataset contained 72,526 records, and included application titles, abstracts and specific aims as well as the study sections that the applications were reviewed in. These applications represented submissions from 42,564 applicants, many of whom submitted multiple applications. The reviewer dataset was retrieved from the NIH Query/Review/Report (QVR) system, a reporting tool for NIH staff, and contained 11,896 unique reviewers. Bibliographic data for the study were obtained from Scopus data (1996-2011, over 25 million records) that were received from Elsevier in summer 2012. Data were parsed from the original XML format into tables holding information about documents, authors, abstracts, etc. from which data needed for this study could be easily retrieved in bulk.

### Identifying Scopus Author-ids for Reviewers

For this study we used Scopus author-ids directly rather than attempting to do our own author disambiguation. Author disambiguation is a current area of research [20] in several fields including scientometrics and machine learning. Author ambiguity occurs in two main ways: multiple authors may have the same name (polysemy), and the same author may have multiple name variations (synonymy, or namesakes). In addition to being an area of research, author disambiguation has attracted commercial interest due to the fact that data vendors have a desire to accurately tag articles to the appropriate authors. Nevertheless, one can assume that it uses information such as co-authorships, title and abstract words, and citation characteristics to make these assignments.

Our experience working with Scopus author-ids enables us to make the following observations [21]. Prolific authors may have more than one author-id (the synonymy problem). However, when this occurs the majority of the author?s publications (and particularly the older ones) are assigned to a single profile. Combining multiple profiles for a single author typically has only a minor effect on total citation counts, and rarely has any effect on the h-index of that author. Even though combining profiles rarely has a large impact on results, we nonetheless identify multiple author-ids for those authors with multiple profiles. There are cases where multiple authors are obviously listed under the same author-id (the polysemy problem). This is the most serious problem with using author profiles because combining publications for multiple authors overstates the expertise and impact of all authors whose profiles are conflated. Cases of polysemy in the Scopus author-ids are typically confined to common author names.

Despite the problems mentioned above, our experience is that use of the Scopus author-ids, after spot-checking and filtering for polysemy, is highly beneficial because the author profiles are much more accurate than if we were to simply search for works of an author using author name and affiliation alone. This is because the Scopus assignment process combines an author?s publications when that author moves from one institution to another or publishes under multiple affiliations, both of which are rather common occurrences for the types of researchers represented in the applicant and reviewer populations.

To characterize SRGs from a reviewer standpoint, Scopus author-ids were identified using the names and affiliations of all reviewers for all meetings held during 2011 and 2012. The total number of unique reviewers in the set was 11,896. Scopus author-ids were determined for 11,779 of these reviewers using a combination of electronic matching and manual review. The process was as follows:

- A list was made of all unique institution names, and shortened strings for each institution were identified (e.g., ?Hopkins? for Johns Hopkins University, ?Michigan? for University of Michigan, etc.) This step was implemented to allow institutional searches to work to a high degree despite institutional name variations in publication data.
- Reviewer names and the shortened institution string were loaded into a database table. The first four characters of the reviewer string were placed in an additional field.
- The entire Scopus author-affiliation data from 2007-2011 was preprocessed and comprised over 45.7 million author-affiliation pairs from 10.56 million unique bibliographic records. The first four characters of the author string were placed in an additional field to be used as a gross lookup feature to match to the four-character reviewer string from the previous step. Institution names, both cleaned (where available, based on previous work), and raw institutional strings were included in this table. This table also included the Scopus author-id and author full name fields.
- The list of reviewer names and the preprocessed Scopus author-affiliation data were joined on the four character name strings to produce a list of possible article matches for each reviewer. This list was further filtered by searching for the shortened institution string (for the reviewer) in the institution name fields (for the Scopus data). These data were then grouped by reviewer name, institution, and Scopus author-id, resulting in counts per name, institution, and author-id triplet.
- Using a pre-computed table containing various data for each Scopus author-id, we added the numbers of Scopus records from 2007-2011 for each author-id. Comparison of this number to the counts from the previous step gives us a good indication as to whether the search strategy identified the majority of the articles associated with each author-id. If the numbers are similar, this indicates that we have likely identified the correct author-id for the reviewer. If the numbers are very dissimilar (ratio ¡ 0.1), it indicates that we should look more closely at that match. There are several reasons that a low ratio could still be a positive match. These include use of name variations by the author that differ from the NIH version of the name, movement of the author from one institution to another, and Unicode character interference.
- Duplicate reviewer name to author-id matches were merged. Duplicates were possible because of the multiple institution fields that were searched for matches to the shortened institution string from the reviewer data.
- From the resulting data file, the list of potential matches was manually inspected. Potential matches were removed if they could not definitively be said to be the same person. Clues that led to removal of a potential match include mismatches on first and middle names or initials, large publication counts (from step 5) that suggested that multiple authors were included in the author-id, low match counts coupled with the reviewer appearing to be at an institution different from the queried institution. Informed judgment was used in all cases.

A total of 10,531 reviewers (88.5%) were matched using this process. Two additional automated runs were made to try to match the remaining reviewers. In the first of these, articles were identified using author plus institution searches against PubMed data rather than using Scopus data. The resulting article list by reviewer was then inserted into the above process repeating steps 5-7, leading to an additional 137 positive matches. In the second of these, the remaining reviewers were run through the full process again but with the institution match removed. This allowed more possible matches for the remaining authors, many of whom were associated with smaller institutions, many of whom for which CSR did not have an institution listed, or for whom the NIH institution listing was wrong. This step led to an additional 612 positive matches. Thus, to this point, 11,280 of our 11,896 reviewers (94.8%) had been matched.

A strictly manual process was used to locate the Scopus author-id for the balance of the reviewer names. The name and institution were input into the online version of Scopus, and the author-ids for obvious matches were manually added to our data. This manual step located author-ids for an additional 499 reviewers. In the course of this combined electronic and manual process of matching reviewers to Scopus author-ids, we identified 91 reviewer names (0.8%) that seemed to be associated with multiple authors of the same name. These reviewer names were excluded from any additional analysis. In addition, no Scopus author-id could be identified using any method for 26 of the reviewers. These were also excluded from additional analysis.

Regarding the issues of polysemy and synonymy in the Scopus author-ids, our data give us some estimates of the rates of occurrence of these phenomena. As mentioned above, 0.8% of the reviewer names seemed to be associated with multiple authors of the same name. The polysemy cases we identified were rather obvious upon inspection. Given that, it is likely that the actual polysemy rate, including less extreme cases, is somewhat higher than this ? perhaps double. Thus, we estimate the polysemy rate within this reviewer pool at 1.5%, with half of those being extreme cases that must be avoided for analysis.

Regarding synonymy, we identified a second Scopus author-id for 870 reviewers (7.3%). All second author-ids identified were associated with at least two articles. The second author-ids for these reviewers accounted for only 7% of their indexed articles and only 3.8% of their total citations. Even though synonymy is a larger issue than polysemy, its effect on our measurements appears to be quite small.

There is one remaining issue regarding author profiles in Scopus. It is possible that some articles not written by the author are incorrectly assigned to the author. Estimation of this quantity would require consultation with authors, and is obviously beyond the scope of this study. However, we assumed that the fraction is relatively low given our experience working with profiles from authors in our own field whose work we know.

### Identifying Scopus Author-ids for Grant Applicants

The process that we used to identify Scopus author-ids for reviewers combined electronic and manual efforts. Given the number of grant applicants for which we needed to identify Scopus author-ids (35,957), we did not use the manual portion of the process that we had used previously. Rather, we designed a scoring system that was intended to mimic the processes that were used to manually inspect potential matches, and used that scoring system to select a single (hopefully best) match for each applicant. The first six steps of the process above were used, and the seventh was replaced by the following: Matches based on single articles were discarded. Matches with obvious mismatches in first name or middle initials were also discarded. Names were not discarded in this step if the full first name was not available in one data source or the other (the NIH names or the Scopus names). For example, ‘J Doe’ and ‘John Doe’ were not considered mismatches in this step. Given that we had a table representing all Scopus author profiles (over 15 million of them), we calculated the number of authors that could have potentially matched a particular name string from the NIH grant data. If this number is low (one, for example), then the chances of the potential match being a true match are high. We also calculated the number of Scopus author-ids identified as potential matches for each applicant. If there are many potential matches, the chance that any of them is a true match decreases. Each potential match was scored using the following formula. The number of papers matching the author and institution strings was squared, and was divided by the total number of papers published by that author profile during the time period matching the search (2007-2011). This is the most dominant feature in our scoring system, and rewards high numbers of matching papers and high recall values for a particular author. The score was augmented by 0.5 if there was an exact match in the first names in the NIH and Scopus data. The score was augmented by 0.25 if only a first initial was used in one data source or the other and that first initial matched the first initial of the full name in the other data source. The score was augmented by 0.5/npot, where npot is the number of potential matches from step 2. The score was also augmented by 0.5/nmatch, where nmatch is the number of matches from step 3. The matches were ranked by score for each applicant, and the top scoring match was kept as the Scopus author-id for the applicant.

Positive matches for identified for 29,800 of the 35,957 applicants for whom we did not already have a Scopus author-id. (There were 6,607 applicants whose author-ids had already been identified because they were also reviewers.) These were subjected to a quick visual screen, and 57 obvious cases of polysemy (several hundreds of papers in the Scopus author profile with very few author name/institution matches) were removed from the match list. The match rate for this automated process was 82.7%, which was slightly less than the match rate obtained for reviewers. However, when applicants who were also reviewers are included in the numbers, Scopus author-ids were identified for 36,350 (85.4%) of the 42,564 applicants. This is still lower than the match rate for reviewers. However, we expect a higher match rate for reviewers because they typically have stronger publishing track records, while some grant applicants may not have a publishing track record that could be matched using our methods.

### Detailed Basemap of Science and Technology

A highly detailed article-based model and map of science have been created using co-citation techniques. Nearly 20 million articles from Scopus (1996-2011) and 2 million U.S. patents comprise this model. A clustering of articles and patents comprises the detailed classification system, and allows portfolios of articles to be compared and visualized. Although patents are a part of this model and map, they are not used in the current study.

The model is formed by taking annual slices of the scientific literature (from Scopus) and U.S. patents (from USPTO data sources), partitioning these literatures into a large number of very small topics, and then linking the resulting annual sets of topics together into time-series based structures. In essence, this model gives a history of science and technology at the level of research problems. One feature of this model is its combination of stable (long-term) and unstable (short-lived) topics, which accurately reflects the actual ebb and flow of work in science and technology at the topic level. This model can be used to analyze existing and hypothetical portfolios in terms of strengths and weaknesses, opportunities and threats, potential collaboration opportunities, topic ages, etc.

The basic, high-level process used to create this model and map of science and technology is as follows: 
- Annual models of science are created from annual slices of Scopus data using co-citation analysis. Creation of each model consists of the following steps.
  - Reference papers are clustered into co-citation clusters using a k50 [22] similarity measure based on the co-citing of these reference papers. Details of this process are available in [23,24].
  - Once these clusters have been created, current year papers are fractionally assigned to the clusters of cited references using a new process that is based on bibliographic coupling [23]. Each cluster thus consists of a set of reference papers and the current papers that build upon them.
- Annual models are then linked into a longitudinal model of science by linking clusters of documents from adjacent years together using overlaps in the cited references belonging to each cluster [23]. Clusters from adjacent years are linked if the cosine overlap between clusters is above a threshold value, which varies between 0.22 and 0.26 depending on year. These linked clusters are referred to as threads. Roughly 40% of the annual clusters are what we call isolates. These are clusters that link neither forward nor backward within the overall model (above a specific threshold), and can thus be thought of as one-year threads. These are research problems that do not have enough momentum to continue into a second year. Isolates are typically among the smallest clusters, while the longest threads are comprised of larger clusters on average. 46% of annual clusters are in threads that last 3 years or longer. The model contains 190,151 article threads of two years or longer.
- The same process is repeated with U.S. patent records. Clusters of documents are created using co-citation analysis, and those clusters are linked into threads. The model contains 29,246 patent threads of two years or longer.
- A visual map was created of all threads aged two years or older from both models (articles from Scopus, patents from the U.S. patent database). BM25 similarity values (equation given below in the Study Section Relatedness section) between pairs of clusters were computed for all pairs of clusters using the titles and abstracts of document in the clusters, and the top-n similarity values for each cluster were used to determine the map layout. Scientific articles and patents are known to use different language; patent abstracts are enriched in legal language and are not always informative as articles. Because of this we kept up to the top-5 links from patent clusters to paper clusters even if their similarity values were lower than patent-patent cluster similarities. This modification allowed the patent clusters to be interspersed with paper clusters to a greater degree than if the modification were not used.
- A system with a smaller number of categories was desirable for the comparison of study section portfolios since a classification system of over 200,000 clusters is perhaps too granular for many types of analysis. Thus, we overlaid the basemap with an 85 × 85 grid that subdivides the map into a smaller number of categories. Given that the mapping process (step 4) places clusters with similar content close together, each grid-based category can be thought of as a science or technology-based specialty. This process results in a set of 4,662 categories that contain scientific articles.

### SRG Relatedness

Choosing the best data to use to determine the relatedness between two different objects is not always straightforward. Co-occurrence in the feature space associated with the objects (e.g., co-term, co-citation) is the most commonly used basis for relatedness [3,25]. As mentioned above, for study sections, we find that we can break the feature space into three main groups, with data types as follows:

- NIH Classification
  - RCDC (Research, Condition, and Disease Categorization) profiles. This measurement is based on the assumption that the relatedness of a pair of study sections scales with overlap (or co-occurrence) in the concepts and terms from the Research, Condition and Disease Categorization (RCDC) in the applications reviewed in each pair. We developed a fingerprint for each study section based on the frequency of occurrence of terms from the RCDC thesaurus in the title, abstract and specific aims of applications reviewed by these study section during the fiscal years of 2011 and 2012. Fingerprints were compared to each other by computing match scores. The scores generated by this process were converted to simple ranks. Thus each study section was compared to each other study section and rank ordered by similarity as defined by the occurrence of terms from the RCDC thesaurus.
- Data associated with reviewers
  - Publications authored by reviewers (REV-P). This metric is based on three assumptions: 1) the topic mix associated with a study section can be represented by the expertise of the reviewers for that study section, 2) the expertise of an author (in this case, a reviewer) can be represented by that author’s publications, and 3) a detailed classification system comprised of publications can be used to determine co-occurrence between two sets of publications. Relatedness between pairs of study sections was calculated after this logic using the following steps. Scopus author-ids were identified for 11,779 of the 11,896 unique reviewers who served in study section meetings during 2011 and 2012. Articles authored by those reviewers were identified and were assigned to the appropriate SRGs, thus giving lists of articles by study section. Each article was weighted by the number of SRGs attended (from one to six) by its author. A total of 642,321 unique study section/article pairs were identified in this step. Using the detailed classification system and basemap of science mentioned above, the number of weighted articles was summed by category for each study section. This step resulted in a matrix of study sections by categories. Relatedness between pairs of SRGs was then calculated as the cosine (or dot product) between the vectors associated with the study sections. A visual representation of relatedness between study sections based on this method is shown in Figure 5. Locations of reviewer publications are shown on the map for each of four study sections from the AARR and BST IRGs. The size of each dot on the map represents the number of publications in that sector of the map. It is very difficult to discern any visual difference between the BSCH and BSPH profiles. Correspondingly, the cosine similarity value for these two SRGs is very high at 0.941. Comparing the bottom two SRGs of Figure 5, similarities and differences can both be seen for the BMBI and NANO study sections, and the corresponding cosine is 0.557. Visual comparison of BSCH and BMBI reveals that there is almost no overlap between the profiles; the resulting cosine is very low at 0.008.
  - Titles and abstracts of publications authored by reviewers (REV-T). This measurement relies on the assumption that two objects are related if the text used to describe them is similar. Textual profiles for each study section were generated from the words in the titles and abstracts of the 642,321 articles mentioned above. Relatedness between pairs of study sections was calculated using the BM25 ranking function. For two documents q and d, BM25 is calculated as:

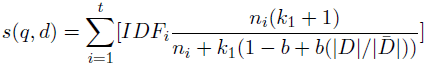

where *n*_*i*_ is the frequency of term *i* in document *d*. The sum is over all *t* terms. Values of 2.0 and 0.75 were used for constants *k*_1_ and b, respectively. In this formulation the text for each study section was treated as if it were a single document. Document length —D— was estimated by summing term frequencies *n*_*i*_ per document. Average document length |*D̄*| is computed over the entire document set. The IDF value for a particular term i was computed as:

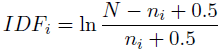

where N is the total number of documents in the dataset and *n*_*i*_ is the number of documents containing term *i*. Each individual term in the summation in the first formula is independent of document q.
  - Cross-citation patterns between reviewers (REV-C). This measurement assumes that two groups of documents are related if documents from one group cite documents in the other group. Using the 642,321 documents authored by reviewers and assigned to SRGs, we calculated the numbers of times papers from each study section cited papers in other SRGs. Self-citations by SRGs were not considered. The remaining count data were used to calculate the fraction of citations that each study section gave to each other study section where the basis for normalization was the sum of citations from the citing study sections.
- Data associated with applications and applicants
  - Text (title, abstract, specific aims) of grant applications (APP-T). This measurement was calculated using the same procedure mentioned above for the REV-T metric. A textual profile was generated for each study section using the titles, abstracts and specific aims sections of the grant applications assigned to it for review. Text from 72,526 grant applications was used for this calculation. BM25 coefficients between all pairs of SRGs were calculated using the textual profiles.
  - Applicant publications (APP-P). This measurement is based on the assumption that grant applicants will write proposals that are similar to their previously published work. The procedure used here was the same as that used for the REV-P metric. The difference was that articles authored by grant applicants were used instead of articles authored by reviewers. Scopus author-ids were identified for 36,306 of the 42,564 unique applicants. The fraction of grant applicants for whom ids were identified is lower than the fraction of reviewers that were positively identified because 1) all matching was done electronically, and 2) we expect that some grant applicants are young scientists with no publication profile. Each article was weighted by the number of individual applications submitted by the applicant to the study section, and the calculation of study section relatedness was done using vector cosines as was done for the REV-P metric.

**Figure 5.**
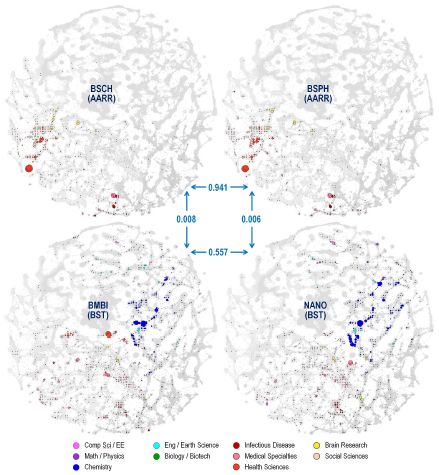
Visual comparison of the locations of reviewer publications for four study sections on the base map of science. Cosine values between pairs of study sections are indicated at the interior of the figure.

For each of the relatedness methods listed above, similarity values were rank ordered (largest similarity first) for each study section. Visualizations of the study section networks were created for each of the six relatedness methods using the following process:

- 174 SRGs were in operation throughout the 2011-2012 fiscal year that framed this study. Another 6 SRGs were in operation during a portion of this time hence a total of 180 SRGs. Each of these is related to nearly every other SRG to some degree. We limited the network visualizations to the strongest relatedness values, and filtered the list of value to keep only the top 3 values per study section.
- A study section network map based on these top 3 similarity values per study section was created using the Kamada-Kawai layout algorithm in Pajek [17].

## Acknowledgments

The authors thank Richard Nakamura for a critical reading of the manuscript and helpful suggestions. The authors also thank Don Tiedemann, Richard Ikeda, and Oliver Morton at NIH for technical assistance and helpful discussions during this study.

